# Simple 3D-Printed Stirred Bioreactor Enhances Retinal Organoid Production Via Improved Oxygenation

**DOI:** 10.1101/2025.06.13.659558

**Authors:** Kyle H. Schwab, Philsang Hwang, Ki Yoon Nam, Zachary Batz, Suja Hiriyanna, Florian Régent, Nicole Y. Morgan, Peter I. Lelkes, Tiansen Li

**Affiliations:** Neurobiology, Neurodegeneration & Repair Laboratory, National Eye Institute, National Institutes of Health, Bethesda, MD, USA; Ocular Gene Therapy Core, National Eye Institute, National Institutes of Health, Bethesda, MD, USA; Department of Bioengineering, Temple University, Philadelphia, PA, USA; Biomedical Engineering and Physical Science Shared Resource, National Institute of Biomedical Imaging and Bioengineering, National Institutes of Health, Bethesda, MD, USA

**Keywords:** Human-Induced Pluripotent Stem Cells, Retinal Organoid, Bioreactor, Oxygenation, Mathematical Modeling

## Abstract

Retinal organoids (ROs), derived from human pluripotent stem cells (hPSCs), simulate *in vivo* development and retinal morphology, providing a platform to study retinal development and diseases. However, current differentiation protocols often yield inconsistent results with substantial cell line and batch variability. These protocols utilize static culture methods that rely on passive oxygen diffusion to reach the vessel bottom, where adherent hPSCs initially differentiate. Static culture is standard for adherent monolayer cells and is presumed suitable for RO differentiation. We questioned this assumption given that, during differentiation, the monolayer hPSCs become highly structured and multi-layered, first as neural rosettes and then as optic vesicles (OVs). We hypothesized that the cellular oxygen consumption rate would exceed the rate of delivery via passive diffusion, particularly to inner regions of emerging OVs. To test this hypothesis, we measured dissolved oxygen concentrations at the vessel bottom and found that within hours of media change, oxygen dropped to < 1 %, a level considered non-physiologically hypoxic, which imperils cell viability. This non-physiological hypoxia caused OV degeneration, hypoxic marker expression, and necrosis. To address this problem, we developed a novel 3D-printed stirred bioreactor (SBR) that maintains physiological oxygen levels between ∼4-6%. This approach significantly improved organoid yield, quality, and reproducibility while being easily adaptable to typical laboratory cell culture workflows. We conclude that non-physiological hypoxia, a previously unappreciated condition, is a limiting factor underlying inconsistent yield and quality in RO production. Physiological oxygenation levels can be restored by the SBR platform, resulting in greater consistency and improved production outcomes.

## INTRODUCTION

Hereditary degenerative retinal diseases, such as retinitis pigmentosa and age-related macular degeneration, are among the leading causes of vision loss in the developed world.^1,2^ Despite significant progress in understanding retinal development and disease pathogenesis using animal models, few therapeutic options have emerged due to disparities between animal and human physiology and the lack of high-fidelity *in vitro* models. Developing accurate *in vitro* human models remains critical to therapeutic advances. Recent studies have demonstrated that human induced pluripotent stem cells (hPSCs) can be differentiated into ROs.^3–6^ ROs replicate key aspects of *in vivo* retinal development and morphology, providing an experimental platform for drug screening, gene therapy, and cell transplantation.^1, 2, 7^ However, current retinal differentiation protocols suffer from technical issues, such as limited yield and inconsistent differentiation efficacy across hPSC lines and production batches, constraining broader applications.^5, 8, 9^

The differentiation of hPSCs into ROs begins with an initial phase of adherent culture lasting 20 – 30 days. This adherent configuration is similar to monolayer culture, which relies on passive diffusion of oxygen and nutrients, and presumes that this method effectively supports hPSC differentiation. However, this assumption ignores a critical difference: differentiating cells increasingly assume complex multi-layered structures, from pseudostratified neural rosettes to optic vesicles (OVs). In fact, cellular architecture significantly impacts oxygen and nutrient availability, presenting a diffusion barrier, as well as hindering removal of waste products.^10^ Additionally, two critical parameters of *in vitro* culture are frequently neglected in RO differentiation protocols: the rate of atmospheric oxygen diffusion through static culture media and cellular oxygen consumption.^10, 11^ These overlooked factors can critically impact differentiation outcomes.^12^

Static culture methods rely on the passive diffusion of oxygen from the air-medium interface downward to adherent cells (Figure 1A). However, passive diffusion alone may be insufficient to support cellular demands. Studies on oxygen diffusion in static culture have shown that this process is inefficient and frequently creates a hypoxic environment if consumption exceeds diffusion.^13–15^ For example, during static culture of early differentiating neural progenitor cells, oxygen consumption can outpace diffusion.^16, 17^ Therefore, hypoxic conditions are likely to prevail under static differentiation of retinal progenitor cells.

**Figure 1.**
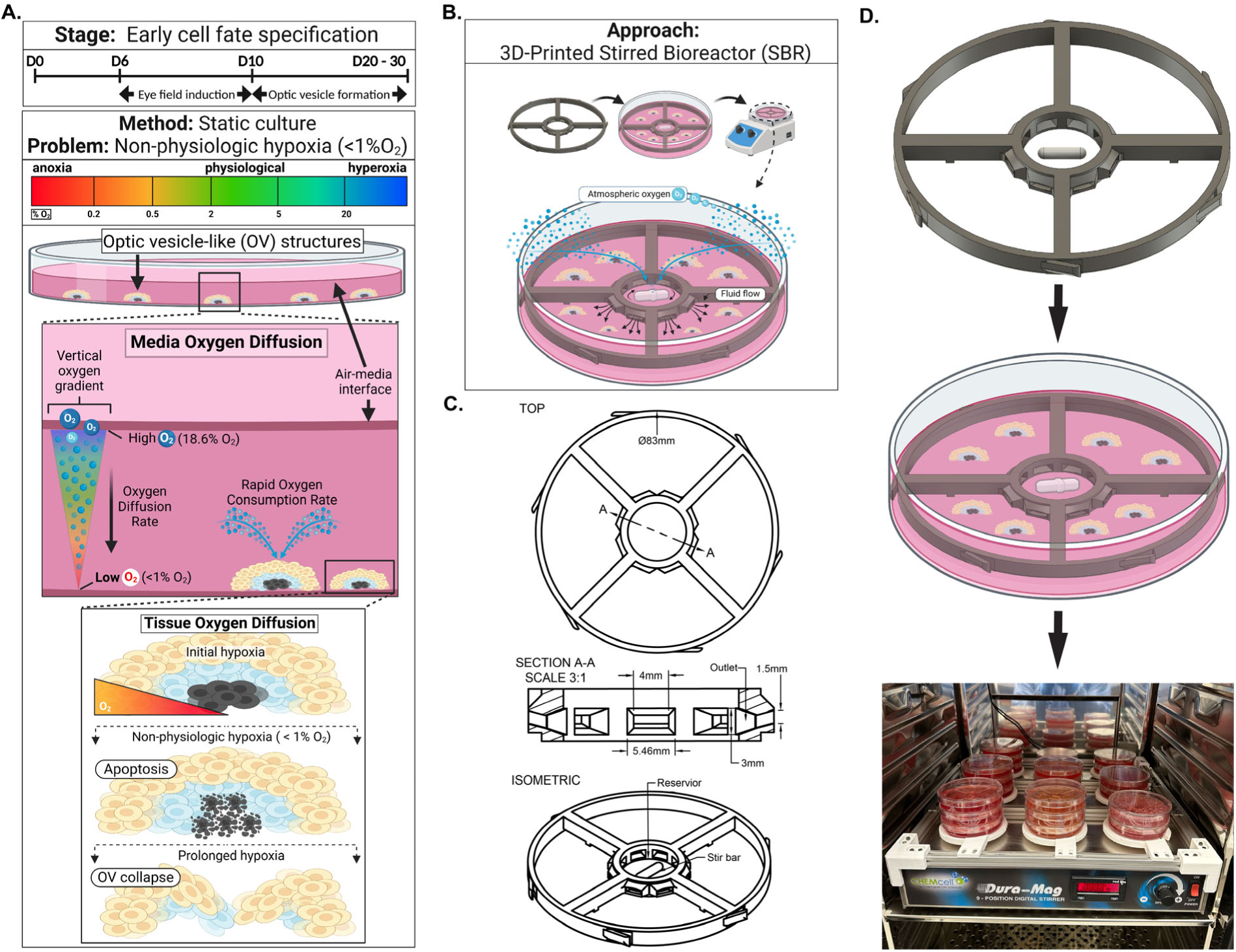
Summary of Problem and Solution, CAD Model of SBR, and Implementation Scheme for Scaled-up Cell Culture Workflows. **<u>A.</u>** Culture stage and illustration of effects of static culture and non-physiologic hypoxia **<u>B.</u>** Approach to overcome non-physiologic hypoxia using SBR **<u>C.</u>** CAD schematic of SBR describing dimensions of geometric features. **<u>D.</u>** Simplified workflow of implemented SBRs during retinal progenitor cell differentiation culture.

Considering that the neural retina is one of the metabolically most active tissues^12,18^ and that *in-vitro* OV-like structures are complex, multi-layered tissues with diffusion limitations^10,18,19^, we posit that passive oxygen diffusion during static culture is insufficient to meet the oxygen demand of nascent and developed OVs. We hypothesize that employing dynamic culture conditions, e.g., media agitation via a stirring mechanism, could improve oxygen diffusion and increase the yield and size of OV-like structures. To test this hypothesis, we designed a novel, 3D-printed stirred bioreactor (SBR) system to enhance oxygen diffusion during early OV formation (Figure 1B-D, Supplemental Video 1). The SBR was designed to be easily implemented in any laboratory setting, using standard 10 cm culture dishes, magnetic stirrer plates, and tissue culture incubators. Combined with our previously described protocol for generating human ROs^8^, the SBR system provides a simple, efficient, and convenient method to increase the yield and size of OVs through improved oxygen delivery and, presumably, more efficient nutrient exchange and waste removal.

## RESULTS

### Differentiation of hPSCs in Static and SBR Culture

The SBR system was validated during initial differentiation using five hPSC lines from day 0 to day 20 (D0 – D20) (Figure 2A, Supplementary Table 1). We followed a protocol in which embryoid body formation was omitted (Hwang et al., manuscript submitted for publication). Briefly, differentiation was initiated in a Petri dish under static conditions (D0 – D6). On D6, the SBR device and magnetic stir bar were introduced and operated at 300 RPM through D20. On D10, hPSC colonies started to show morphologies akin to neuroectoderm formation, establishing eye fields composed of pseudostratified neuroepithelial layers that formed crescent structures.^19^ Within these structures, neural stem cells organized into rosettes, exhibiting radial morphology within 10–15 days (Figure 2B, black arrows). Notably, distinct differences emerged between SBR and static cultures at D15, where rosettes appeared larger and more numerous, with more clearly defined margins and lumens under SBR conditions. Continued development of the neuroepithelia within the rosettes led to the generation of OV-like structures in both static and SBR conditions (D15 - D17, white arrows). In static cultures, many of the OV-like structures were often transient, appearing to undergo a degenerative process and disappeared (D21, Static, white arrowhead). The viable OV-like structures that remained rarely developed clear exterior borders and well-defined lumens. In contrast, the SBR-cultured OVs exhibited thick, well-defined neuroepithelial layer outlines (outer margin and lumen), as shown in phase-contrast images (D21, SBR, black arrowhead). Cell fate specification was confirmed using markers for retinal progenitor cells CHX10 and apical tight junction marker ZO1, which are established markers of the presumptive neural retina and apical junctional complexes of neural epithelia, respectively (Figure 2C). Fate specification were similar between SBR and static conditions. However, under static conditions, CHX10^+^ cells (green) exhibited diffuse, flat patterning. In contrast, SBR-cultured OVs showed peripheral CHX10^+^ cell localization, suggesting a more three-dimensional architecture with luminal topography. Further structural differences can be seen in peripheral ZO1^+^ cell patterning (red), with a planar monolayer-like layer in static-cultured OVs compared to a thin, compact layer in SBR culture. structures, neural stem cells organized into rosettes, exhibiting radial morphology within 10–15 days (Figure 2B, black arrows). Notably, distinct differences emerged between SBR and static cultures at D15, where, under SBR conditions, rosettes appeared larger and more numerous, with more clearly defined margins and lumens. Continued development of the neuroepithelia within the rosettes led to the generation of OV-like structures in both static and SBR conditions (D15 - D17, white arrows). In static cultures, many of the OV-like structures were often transient, appearing to undergo a degenerative process and disappeared (D21, Static, white arrowhead). The viable OV-like structures that remained rarely developed clear exterior borders and well-defined lumens. In contrast, the SBR-cultured OVs exhibited thick, well-defined neuroepithelial layer outlines (outer margin and lumen), as shown in phase-contrast images (D21, SBR, black arrowhead). Cell fate specification was confirmed using markers for retinal progenitor cells CHX10 and apical tight junction marker ZO1, which are established markers of the presumptive neural retina and apical junctional complexes of neural epithelia, respectively (Figure 2C). Fate specification was similar between SBR and static conditions. However, under static conditions, CHX10^+^ cells (green) exhibited diffuse, flat patterning. In contrast, SBR-cultured OVs showed peripheral CHX10^+^ cell localization, suggesting a more three-dimensional architecture with luminal topography. Further structural differences can be seen in peripheral ZO1^+^ cell patterning (red), with a planar monolayer-like layer in static-cultured OVs compared to a thin, compact layer in SBR culture.

**Figure 2.**
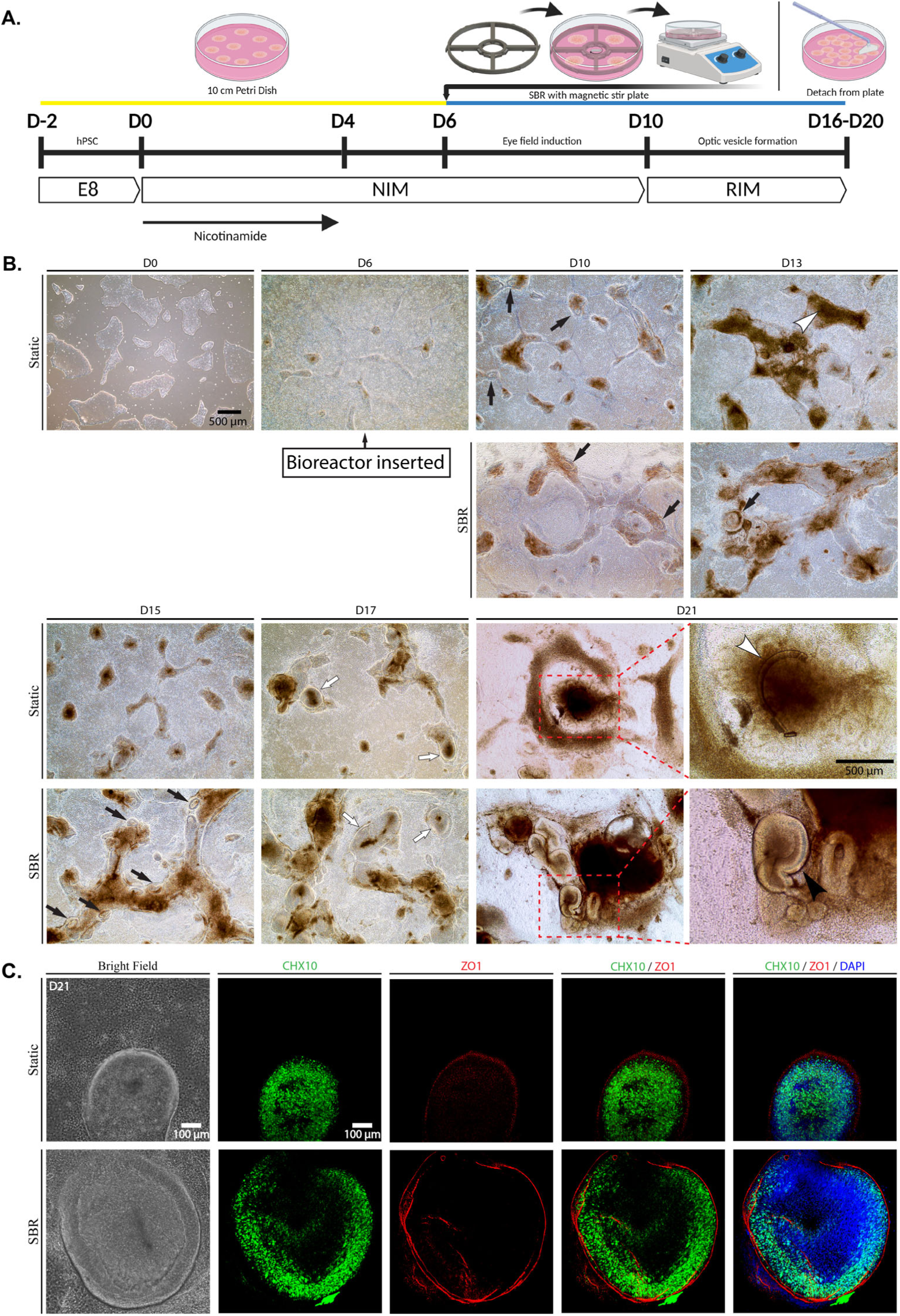
Description of SBR protocol used for differentiating ROs. **<u>A.</u>** Schematic representation of differentiation using SBR. hPSC: human pluripotent stem cells; SBR: stirred bioreactor; NIM: neural-induction media; RIM: retinal-induction media. **<u>B.</u>** Representative bright-field images showing differentiation progression of cell line hPSC1. **<u>C.</u>** Immunohistochemistry analysis of adherent OVs (hPSC1) using antibodies against markers for retinal progenitor cells or bipolar cells (CHX10, green), tight junction protein zonula occludens (ZO1, red). Nuclei were stained with 4′,6-diamidino-2-phenylindole (DAPI, blue). White arrows with black outline point to dashed red (Static condition) and blue (SBR condition) outlines, highlighting differences in outer neural epithelial contours.

### SBR Increased Yield and Size of Retinal Organoids

From D20 - D25, adherent OVs were mechanically detached and transitioned to suspension culture. After 48 hours, ROs formed under both static and SBR conditions, identifiable by a distinct phase-bright neuroepithelial rim (Figure 3A). Compared to static conditions, SBR culture generated a ∼2 to 4-fold higher yield of ROs (Figure 3B, Supplementary Table 3), with a ∼2-fold increase in cross-sectional areas (Figure 3C, Supplementary Table 3). Results were consistent across differentiation protocols (adherent hPSC colony and embryoid body) (Figure 3B–C). Chx10^+^ cell layers in SBR-cultured ROs was significantly thicker compared to static conditions (Supplmentary Figure 1B).

**Figure 3.**
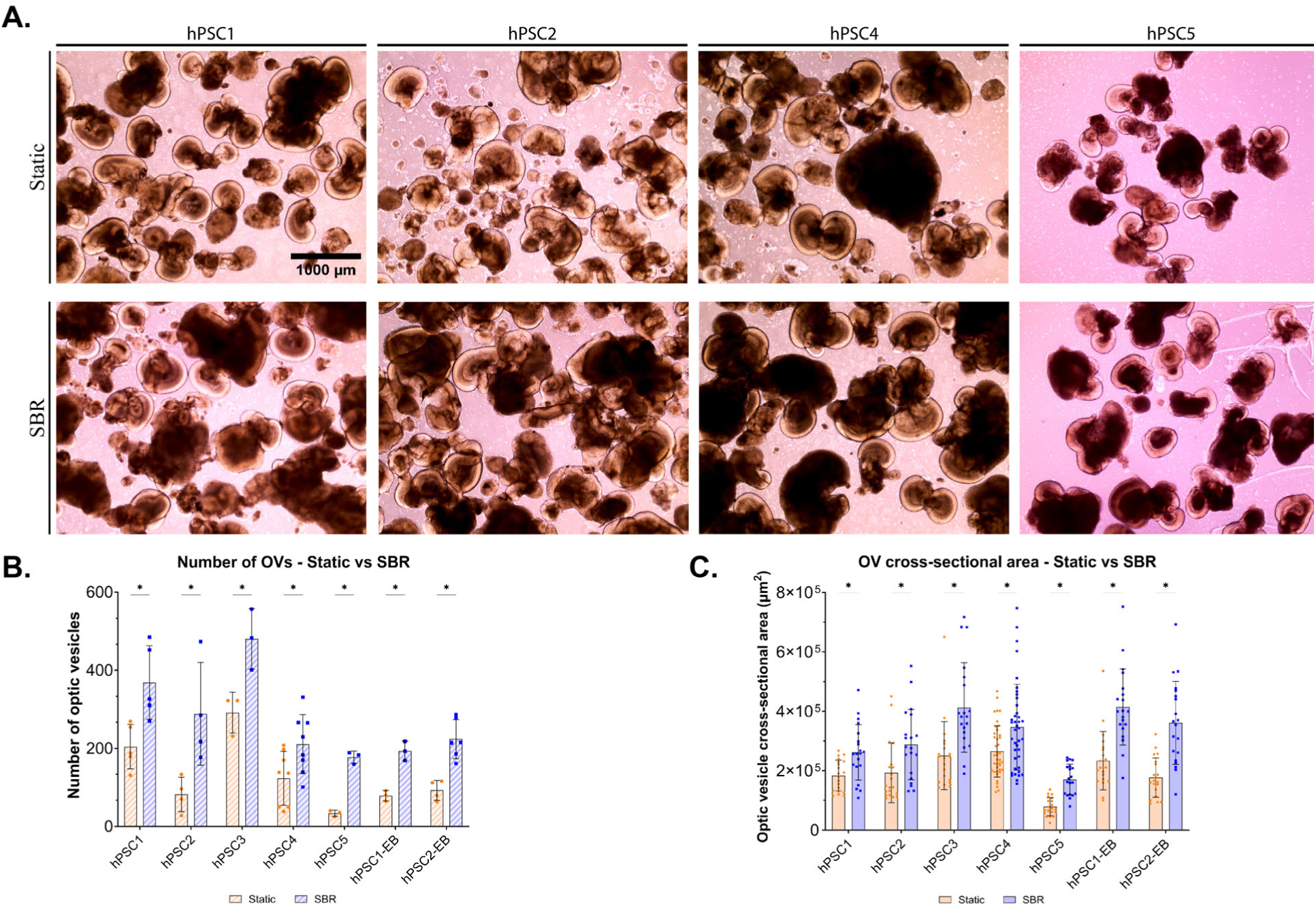
SBR improves yield and size of ROs. **<u>A.</u>** Representative bright-field images showing ROs derived hPSCs from static and SBR conditions. **<u>B.</u>** Quantification of the number of ROs produced by static and SBR conditions. EB: embryoid body. The bar charts summarized data from three (hPSC3, hPSC5, hPSC1-EB, hPSC2-EB) and six (hPSC1, hPSC2, hPSC5) independent experiments. **<u>C.</u>** Quantification of the cross-sectional area (µm^2^) produced by static and SBR conditions. The bar charts summarized data from twenty measurements. All experiments used four different hPSC lines and presented as mean ± standard deviation (p-values shown in Supplementary Table 3).

### SBR Platform Achieved Physiological Oxygen Concentration in Retinal Organoid Culture

Measurement of available oxygen in static cultures, following media change, indicated a steady reduction in oxygen concentrations, with hypoxic conditions occurring more rapidly later in OV development, in line with the increase in cell density and metabolic demands (Figure 4A, Supplemental Figure 1A). Between days 11–13, oxygen levels neared non-physiologic hypoxia (≤1% O₂) about 40 hours after media change (Figure 4B). Oxygen depletion below 1% occurred progressively faster at D13– 15 (17.5 hours), D15–17 (12.5 hours), and D17–19 (6.2 hours). Increasingly complex neuroepithelial structures coincided with oxygen levels falling below physiological thresholds, although total cell counts appeared to have plateaued after D15 (Supplemental Figure 1A). Comparison of oxygen consumption profiles between static and SBR cultures averaged between D11–D19, clearly indicated rapid progression to hypoxia (<1% O₂ within 6 hours) under static conditions (Figure 4C). In contrast, SBR cultures exhibited only a limited reduction in oxygen, reaching a level of equilibrium above ∼4%. To illustrate the robust and consistent effect of SBR operation, oxygen levels were measured under alternating ON-OFF SBR operation and showed highly reproducible peak and trough levels. (Figure 4D).

**Figure 4.**
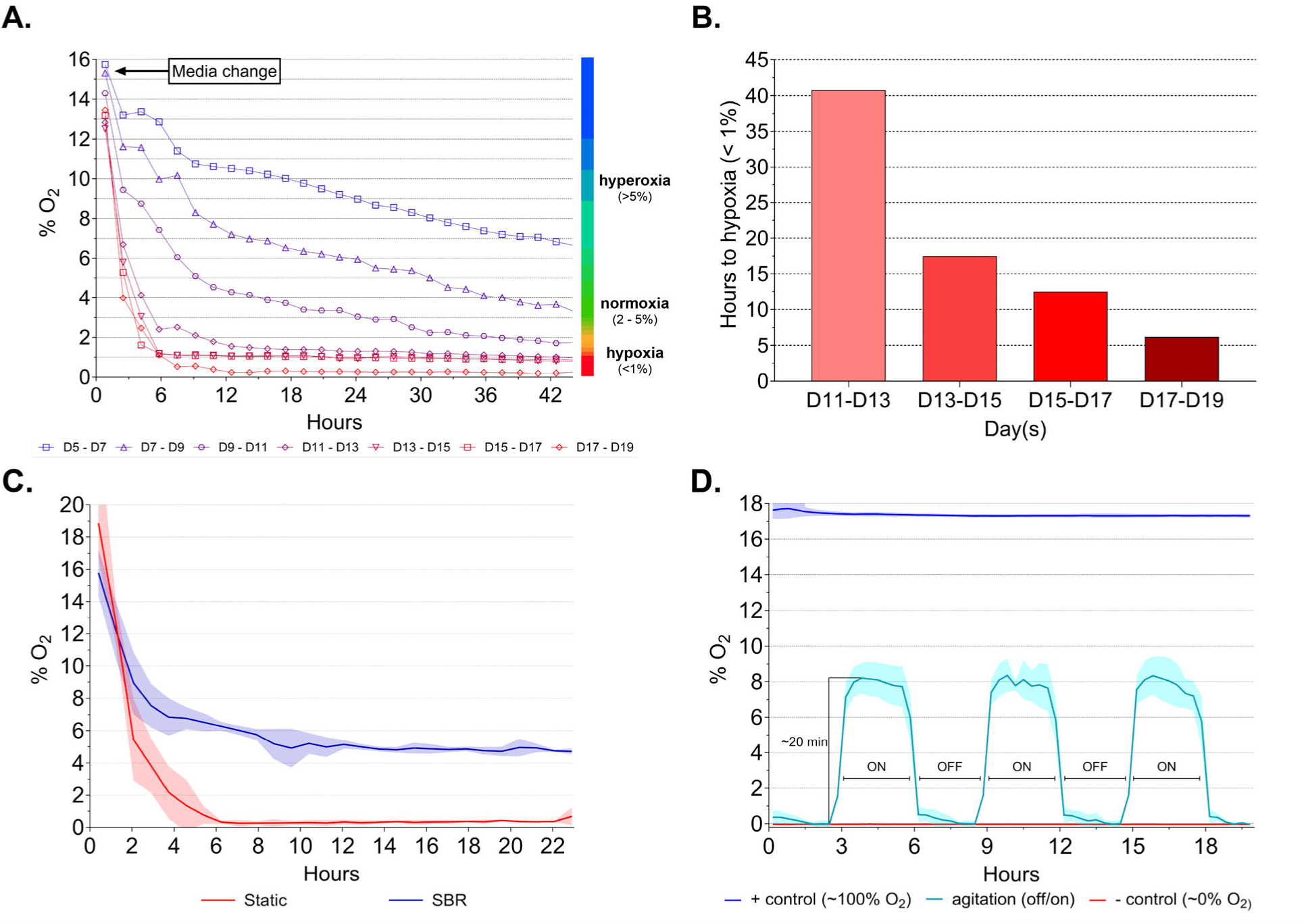
Oxygen (%O_2_) consumption during differentiation of ROs. **<u>A.</u>** Quantification %O_2_ during density-dependent oxygen consumption over 44 h period from D5 – D19 of static differentiation of hPSC-derived ROs (hPSC1). **<u>B.</u>** Hours until available O_2_ drops below 1% in static culture. **<u>C.</u>** Quantification of % O_2_ using continuous agitation over 20 hours. **<u>D.</u>** Quantification of %O_2_ using intermittent agitation with 3 hours stirred (ON) and 3 hours static (OFF), demonstrating cyclic oxygenation of culture media using SBR.

### Deployment of SBR Reduced Hypoxia and Necrosis in Developing Optic Vesicles

Pimonidazole staining is an assay that can reveal severe hypoxia in cells. Under static conditions, OVs showed an increase in pimonidazole-positive cells throughout the neuroepithelial layer compared to SBR-cultured OVs (Figure 5A). A significantly higher mean fluorescence intensity of pimonidazole staining (A.U. - arbitrary units) was observed under static conditions compared to SBR (Figure 5B, Supplementary Table 4). These results confirmed that under static conditions, OVs were exposed to significantly lower oxygen concentrations compared to SBR culture. Consequently, the impact of hypoxia became evident through apoptosis assays. TUNEL assays, which detect cell death, indicated a higher number of apoptotic cells in static-cultured OVs compared to OVs maintained in SBR (Figure 5C, D, and Supplementary Table 4). Overall, static-cultured OVs exhibited a significantly greater number of apoptotic cells (2.6 to 3.5-fold increase) compared to SBR conditions.

**Figure 5.**
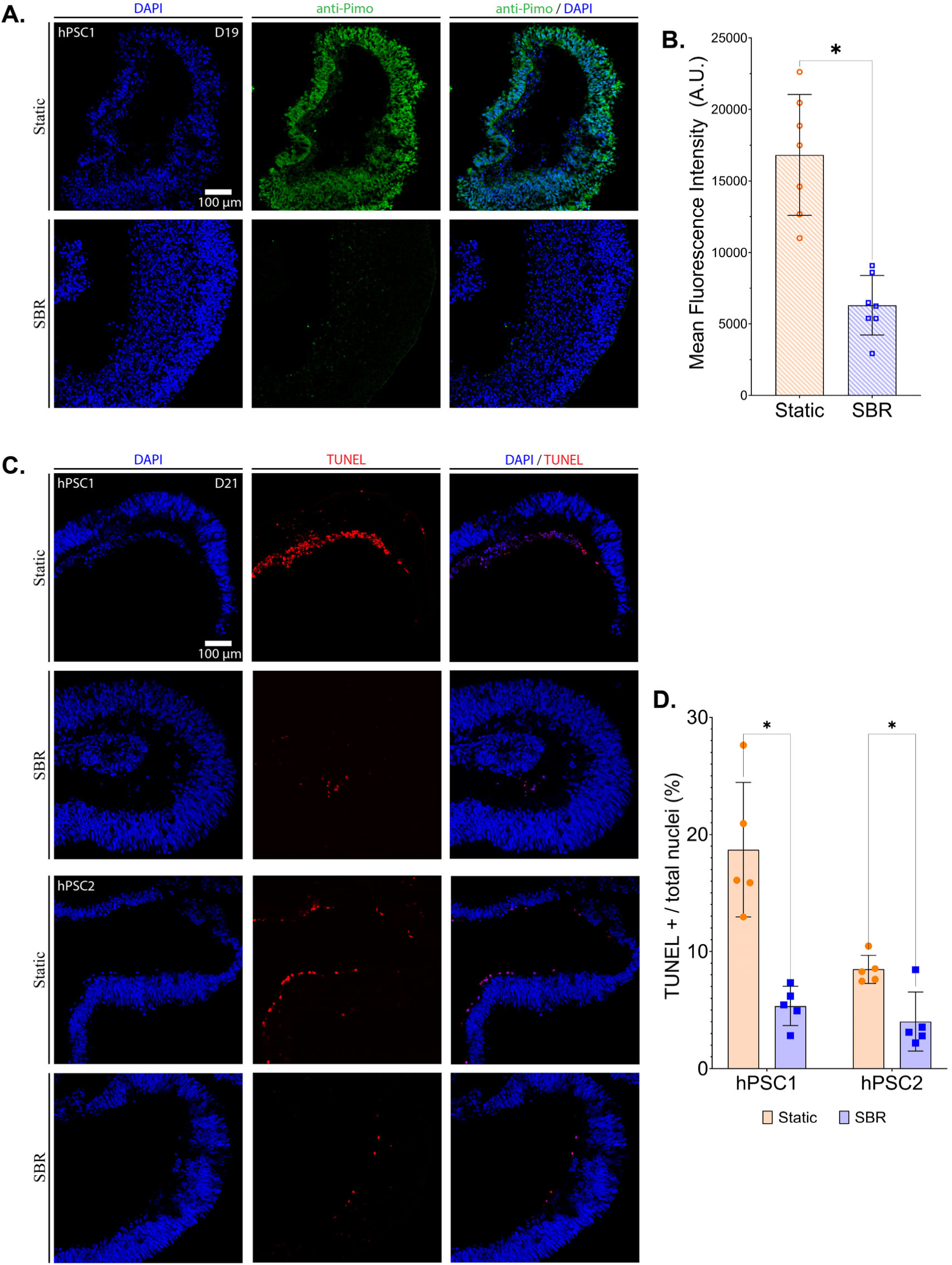
Detection of Hypoxia Gradients and Apoptosis. **<u>A.</u>** Immunohistochemistry analysis of hPSC-derived OV cryo-sections using antibodies against markers for hypoxic cells (antiPimo, green) with nuclei stained with 4′,6-diamidino-2-phenylindole (DAPI, blue). **<u>B.</u>** Quantification of mean fluorescent intensity (A.U.) of hypoxic cells within the OV. The bar charts summarize data from six measurements. **<u>C.</u>** Immunohistochemistry analysis of hPSC-derived OV cryo-sections using antibodies against markers for cells undergoing DNA fragmentation (apoptosis marker) (TUNEL, red) with nuclei stained with 4′,6-diamidino-2-phenylindole (DAPI, blue). **<u>D.</u>** Quantification of TUNEL-positive cells over total nuclei (percentage) compared between OVs in static and SBR conditions. The bar charts summarize data from five measurements. All data are presented as mean ± standard deviation (p-values shown in Supplementary Table 3).

### Physical and Numerical Modelling of SBR Fluid Dynamics

To better understand the fluid flow pattern generated by SBR operation, a physical flow model was created using fluorescent microspheres (Figure 6A, Supplemental Video 2). The trajectories of 63 individual spheres were tracked over a 100-ms exposure period, yielding an average particle velocity of 6.60 ± 2.93 mm/s. These data were compared to time-averaged (t = 0 – 10 s) fluid velocity vector profiles and magnitude contours derived from computational fluid dynamics (CFD) simulations. The CFD vector profiles revealed a recirculating flow pattern in which fluid exited through the outlet ports toward the tissue culture area and was drawn back into the central reservoir by the pressure gradient created by stirring (Figure 6B, Supplemental Video 3). Within the sample observation window (Figure 6B, blue arrow), simulated velocity magnitudes ranged from 2.6 to 8.3 mm/s, closely matching physical measurements, suggesting that the CFD-predicted shear stress values accurately represent the physical conditions. Shear stress contours in the XY-plane (Figure 6C, top-down view, 0.125 mm above the culture dish bottom) and in the ZY-plane (Figure 6D, cross-sectional view) indicated shear stress values ranging from 0 to 0.002 Pa within the OV culture region. Notably, the highest shear stress values were localized near the outlet ports and under support pillars, suggesting that OVs in this region may be exposed to the upper limits of the shear stress range (Figure 6C, D). Importantly, the shear stress experienced in these areas is well below the levels that might damage neuroprogenitor cells in culture, as reported in the literature.^19, 20^

**Figure 6.**
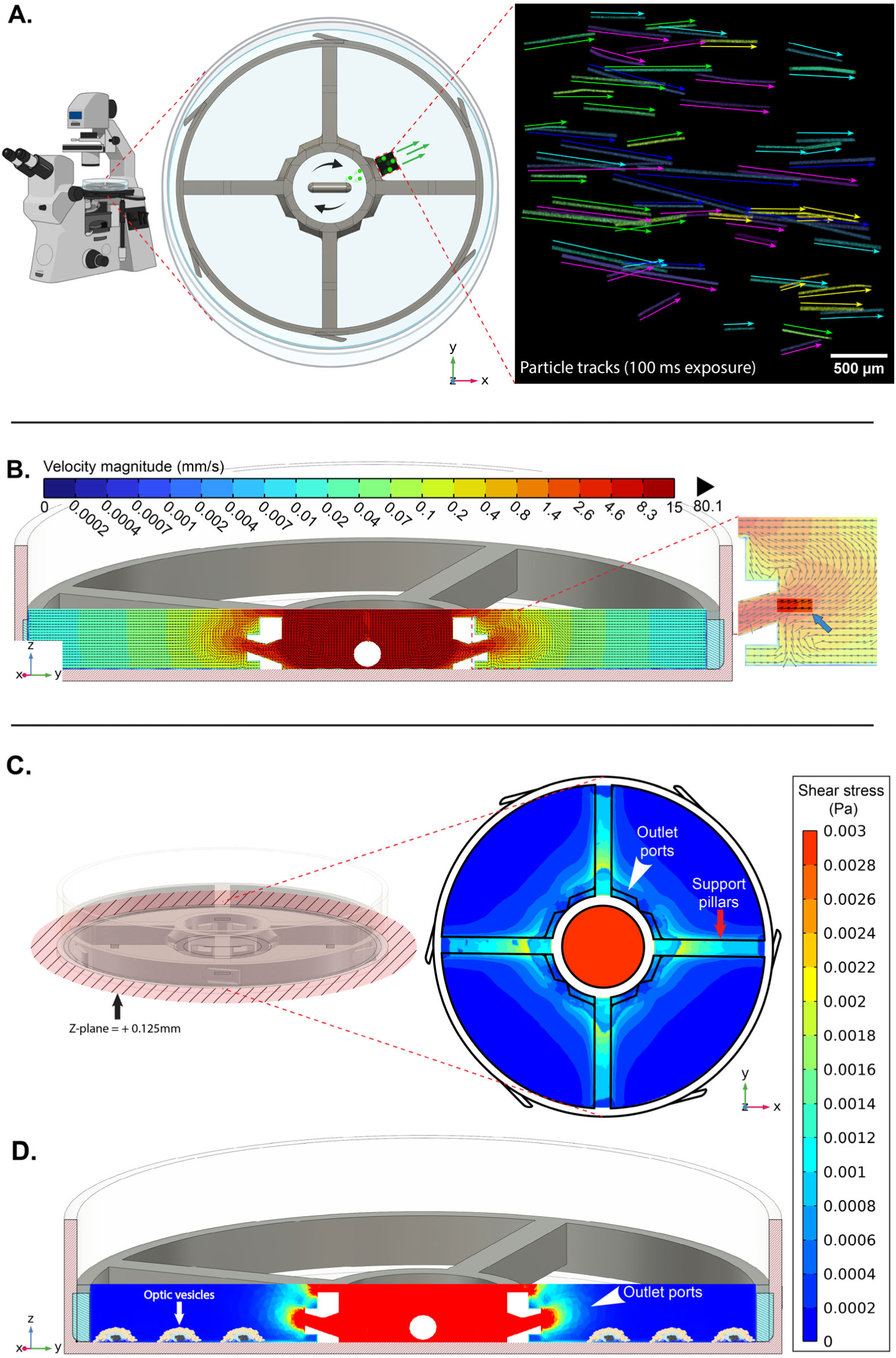
Computational Fluid Dynamics Analysis. **A.** Simplified PIV setup and captured particle tracks at XY-plane of Z = +2.5 mm **<u>B.</u>** CFD fluid flow vector profile of velocity magnitude on ZY-cross-section. **<u>C.</u>** Averaged shear stress contours (0-10 seconds) at bottom (Z = +0.125mm) of Petri dish. **<u>D.</u>** Averaged shear stress contours on ZY-cross-section (0-10 seconds).

### RNAseq Transcriptome Analysis of SBR vs. Static Cultured Retinal Organoids

SBR induces media flow sweeping over cell surfaces, thereby introducing a degree of shear stress (between 0 – 0.002 Pa) upon developing ROs, which may impact stem cell differentiation.^19^ To assess if fluid movement altered cell fate specification and RO development, we performed gene expression profiling of organoids in static and SBR cultures from two hPSC lines.

We first assessed global expression patterns using principal component analysis (PCA). Culture conditions (SBR or static) are captured in principal component 3 (PC3; Figure 7A) and account for 10% of the variation observed in expression across our samples (Figure 7B). Notably, variation in gene expression is more strongly associated with variation associated with patient line (25%, PC2; Figure 7A, 7B) as well as stochastic heterogeneity between samples (32%, PC1; Figure 7A, 7B). Given heterogeneity between cell lines, we investigated expression changes linked to culture differences within individual patient lines. We identified 235 and 39 genes differentially expressed in hPSC1 and hPSC2, respectively, including a total of 18 genes that were shared in both lines (Figure 7C).

**Figure 7.**
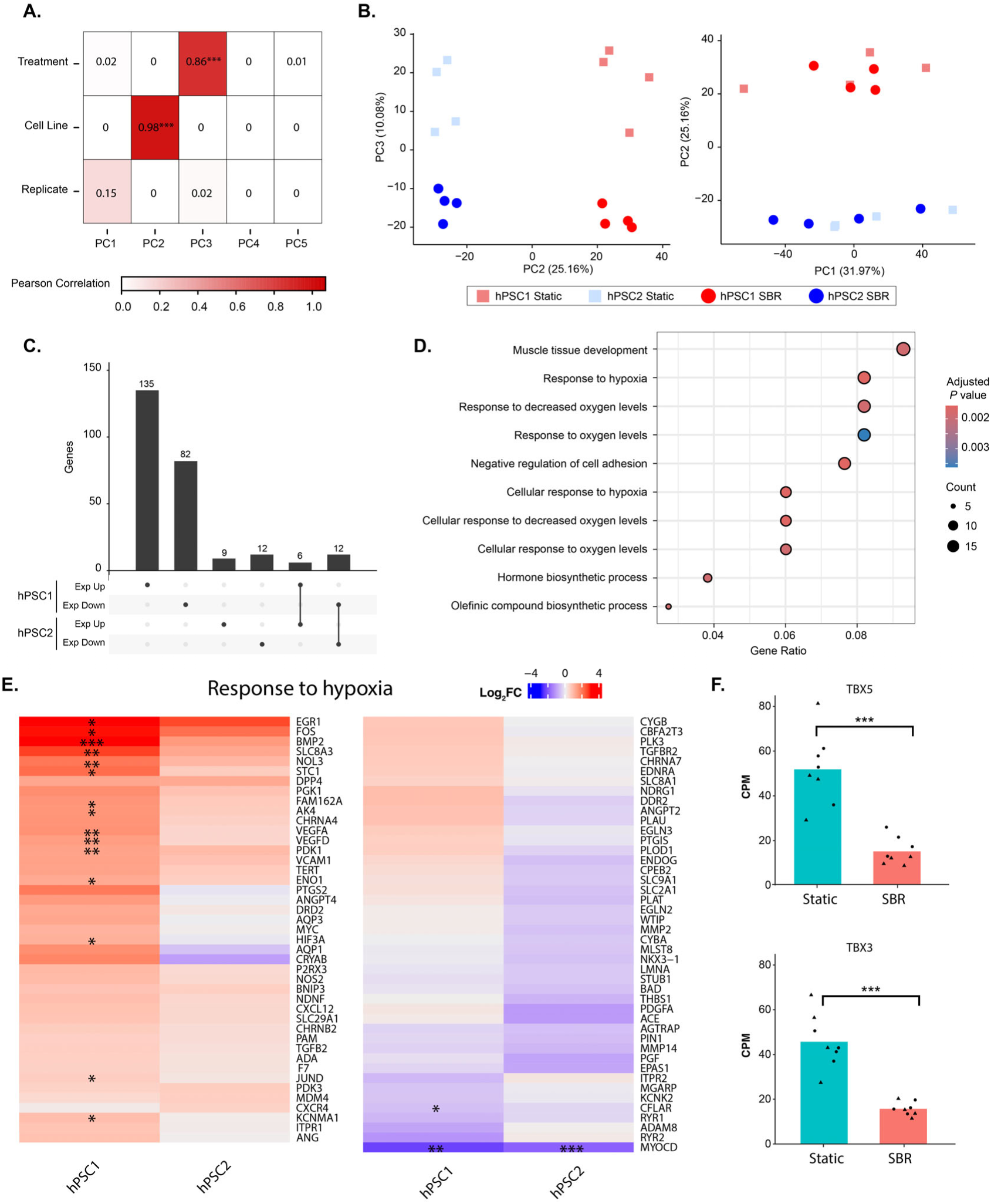
Gene Expression Profiling. **<u>A.</u>** Pearson correlation between experimental factors and principal components derived from transcriptome data. *** indicates a significant correlation at the p < 0.001 level. **<u>B.</u>** Principal components summarizing global transcriptional patterns on PC1 and PC2 (left plot); PC2 and PC3 (right plot). Red and blue points indicate hPSC1 and hPSC2, respectively. Square and round points indicate static and SBR culture conditions, respectively. **<u>C.</u>** Number of differentially expressed genes split by direction of expression change and classified based on their presence or absence in either cell line. **<u>D.</u>** GO terms enriched (p < 0.01) among genes that were differentially expressed in only the hPSC2 cell line. Point size indicates the number of genes identified in a given GO term, color scale indicates the p value associated with the enrichment of each term. **<u>E.</u>** Gene expression changes in SBR vs static conditions identified as part of the “Response to Hypoxia” GO term (GO: 0001666). * indicates p < 0.05; ** p < 0.01; *** p < 0.001 within a specific cell line. **<u>F.</u>** Gene expression in counts per million (CPM) for TBX3 and TBX5 genes. Each point indicates the CPM obtained from a single sample, Bars indicate the mean of all samples. *** indicates FDR < 0.001.

Genes that were differentially expressed in only hPSC2 were enriched for melanin biosynthesis due to downregulation of PMEL and TYRP1 (both associated with RPE), while genes differentially expressed in only hPSC1 were primarily enriched for terms associated with hypoxia (Figure 7D). Under the GO category “*Response to Hypoxia*”, SBR-cultured hPSC1 ROs exhibited higher expression of oxygen-responsive genes compared to static controls (Figure 7E). Enrichment analysis of genes differentially expressed in both cell lines identified GO terms related to cardiac and smooth muscle development due to reduced expression of both TBX3 and TBX5 (Figure 7F). Overall, we saw no significant differential expression in neural retinal cell fate markers and no indication of altered vascular endothelial development. Therefore, fluid flow due to SBR did not introduce any significant or unwanted impact on RO differentiation.

## DISCUSSION

Oxygen availability is fundamental to cell survival, differentiation, and tissue development. During early embryogenesis, non-vascularized tissues encounter physiological hypoxia (1–6% O₂, 8–45 mmHg), particularly in the developing central nervous system.^13, 21^ Levels below 1% represent severe, non-physiological hypoxia detrimental to neural development. Physiologic normoxia (2–5% O₂) is essential for proliferation and tissue homeostasis *in utero*, influencing metabolic and developmental trajectories in complex tissues such as the retina.^16–20^ The mature retina’s high metabolic demands underscore the necessity of precise oxygen regulation.^22, 23^ Replicating this normoxia *in vitro* remains challenging. Static culture systems, the predominant method for RO generation, rely on passive oxygen diffusion, which is often insufficient due to oxygen solubility limits and substantial diffusion distances (millimeters) compared to the ∼30 µm capillary proximity *in vivo*.^23^ This discrepancy results in hypoxic regions within culture media and tissues, compromising OV yields, size, and differentiation outcomes. Severe hypoxia significantly contributes to inconsistent outcomes in static adherent cultures, as evidenced by many emerging OV-like structures that undergo deterioration and fail to survive.^24^

Adoption of the SBR platform, providing physiological oxygen levels, mitigates this deterioration. While oxygen quantification was emphasized, enhanced nutrient delivery and waste removal likely further improved outcomes. Previous bioreactor studies focused on later-stage, free-floating cultures^25, 26^, whereas our study was the first to demonstrate benefits during the initial adherent differentiation stage.

Static conditions, although typical for adherent cultures, inadequately support RO differentiation due to rapidly decreasing oxygen levels below physiological limits (<1% O₂) after media replacement. The pseudostratified structure of neural rosettes exacerbates oxygen limitation, contrasting monolayers with uniform oxygen access. Neurospheres similarly experience core necrosis due to hypoxia.^27^ We often observed cell death in OV core regions under static conditions likely resulting from severe hypoxia-induced HIF-1α activation, triggering apoptotic pathways and impairing differentiation into retinal lineages.^28, 29^ Under prolonged hypoxic conditions, HIF-1α upregulates pro-apoptotic factors, leading to significant cell death, particularly in OV core regions where oxygen diffusion is most limited.^10, 18^ Under extended oxygen deprivation, cells can also upregulate hypoxia-resistant genes, resulting in impaired proliferation, oxidative damage, and DNA fragmentation.^13–15^ For RO cultures, these findings underline the limitations of static culture systems. In theory, reducing media volume and depth could improve oxygen transfer to the bottom of the culture dish.^11, 12^ However, this is not practical given the high metabolic demand for nutrients. Consequently, the formation of hypoxic regions represents a critical bottleneck, not just for survival but for the overall quality and reproducibility of the organoids produced. Therefore, overcoming oxygen limitations requires a culture configuration that meets the increased oxygen and metabolic demands of differentiating cells.

To overcome the oxygen limitations of static cultures, we developed a 3D-printed stirred bioreactor (SBR) (Figure 1B), employing gentle agitation to maintain physiologic oxygen levels (2–5% O₂), avoiding extreme hypoxia. Continuous measurements confirmed sustained physiological oxygenation even under high metabolic demand.^13, 30^ Consequently, SBR conditions produced significantly larger organoids and higher yields compared to static conditions. Immunohistochemical analysis revealed that the neuroepithelial layers in SBR-cultured OVs were more defined and contained significantly fewer hypoxic (pimonidazole-positive) cells, underscoring the efficacy of the oxygen-rich microenvironment.^13^ Continual oxygen delivery and maintenance of physiologic oxygen levels not only improved OV survival but also supported uninterrupted morphogenesis, yielding larger and more structurally complex ROs.^30^ By satisfying the metabolic demands of differentiating cells, the SBR may be able to remove one of the major barriers to scalable and reproducible RO production.

In addition to ascertaining oxygen supply, the SBR addresses mass transport, another critical limitation of static cultures. Adequate nutrient delivery and waste removal are essential for high-density tissue viability. Static systems rely on inherently slow passive diffusion, frequently causing nutrient gradients and waste accumulation that impair tissue development.^10^ The convective flow introduced by SBR stirring ensures improved nutrient distribution and efficient removal of harmful metabolic byproducts.^18^ Although we did not directly measure nutrient or waste diffusion in this study, it is likely that the SBR’s media agitation minimized any such gradients, consistent with foundational studies on diffusion kinetics and dynamic culture methods.^10, 31^ Improved mass transport may therefore partly explain the enhanced survival and growth observed in SBR-cultured organoids.

While delivering marked improvement to the culture conditions, SBR also introduces shear forces that can influence cellular behavior. Shear stress can activate mechano-transduction pathways that regulate cell fate and proliferation.^32, 33^ Computational fluid dynamics simulations confirmed that the average shear stress generated in SBR culture (0 – 0.002 Pa) remains within tolerable limits for neural progenitor cells. These values align with previous reports that shear stress in the 10^-4^ - 10^-3^ Pa range enhances cell proliferation and promotes fate specification.^32, 19, 34^ Interestingly, OVs near the outlet ports and under the pillars, where shear stress is highest (0.001 - 0.002 Pa), tend to grow larger, suggesting a possible positive correlation between fluid flow rate and growth or OV development.

In our study, the amount of shear stress experienced by SBR cultured cells was well tolerated, with no substantial changes in cell fate specification as shown by gene expression profiling from RNAseq analyses. Variation in gene expression is more strongly associated with variation associated with patient lines, as well as stochastic heterogeneity between samples, suggesting that under our conditions the low-level shear stress from media flow did not significantly influence cell differentiation pathways.

As mentioned above, under the GO category “*Response to hypoxia*”, SBR-cultured hPSC1 ROs exhibited higher expression of oxygen-responsive genes. Specifically, a set of immediate-early transcriptional regulators (e.g., EGR1, FOS), growth factors and signaling molecules (BMP2), and a broad array of classical hypoxia-inducible targets involved in glycolytic metabolism and survival (PGK1, ENO1, PDK3, AK4, STC1). Pro-angiogenic factors VEGFA and VEGFD, often correlated with hypoxic responses, were also upregulated in SBR cultures. These findings would seem paradoxical at first glance. Widespread upregulation of hypoxia response genes was an unexpected result and may be linked to the collection method employed. Specifically, samples for RNAseq were collected two days after conversion of the cultures from adherent to free-floating configuration. This method likely exposes static organoids to increased oxygenation but reduces oxygenation levels for SBR-cultured organoids. We note that only one of the two cell lines exhibited this response. Furthermore, we found no significant change in a key hypoxia response gene HIF-1α, that is dynamically regulated on a time scale of minutes to hours.^35,36^ RNAseq analysis performed on samples taken during the adherent phase of culture should capture the actual state of hypoxia and normoxia, respectively. Additionally, enrichment analysis of genes differentially expressed in both cell lines identified GO terms related to cardiac and smooth muscle development due to reduced expression of both TBX3 and TBX5. These two genes are closely linked members of the T-box transcription factor family involved in vasculogenesis in the heart. TBX3 has been shown to play a similar role in retinal vascularization as well.^36,37^ The reduced expression of transcription factors in both cell lines may be due to reduced demand for vascular development in highly oxygenated SBR cultures.

In summary, our study identified severe hypoxia during the initial adherent phase of RO differentiation as a critical bottleneck and introduced a dynamic culture system overcoming key static method limitations. The novel SBR platform, adaptable to various laboratory settings, significantly increases oxygen levels and improves mass transport, enhancing RO yield, size, viability, reproducibility, and scalability. The versatile SBR design could be extended to other organoid systems (e.g., cerebral, hepatic, cardiac), providing a simple solution to improve oxygenation and nutrient uniformity. For RO cultures specifically, enhanced oxygen delivery relieves metabolic constraints, supporting consistent organotypic differentiation and addressing major limitations inherent to static culture methods.

## Supporting information

SBR 3D Model

Supplementary Video 2 - Velocity Cross section

Supplementary Video 3 - Shear Stress

## Acknowledgments

We thank Dr. Robert Fariss of the NEI Imaging Core for help with confocal microscopy. Dr. Milton English for performing RNAseq data collection. This research was supported [in part] by the Intramural Research Program of the NIH, National Eye Institute. This research was supported by the National Eye Institute (EY000490), NIH.

## Abbreviations

hPSCs: Human induced pluripotent stem cells
RO: Retinal organoid
OV: Optic vesicle
SBR: Stirred Bioreactor

## METHODS

### Design, Fabrication, and Implementation of SBRs

The SBR was designed using computer-aided design software (Fusion360, Autodesk). As seen in Figure 1C, the design of the SBR 3D model incorporated critical features such as a central reservoir, angled fluid flow outlet ports, and Petri dish retention tabs. As a critical design criterion, this device had to fit into standard 100 x 20 mm Petri dishes and accommodate 12 x 3 mm magnetic stir bars within the central reservoir. Stirring in the central reservoir was designed to generate a laminar flow through outlet ports angled upwards, driving the fluid movement toward the air-liquid surface. Stirred bioreactors were fabricated using a biocompatible resin (BioMed Clear, Formlabs, USA) and an SLA 3D-Printer (Form 3b+, Formlabs, USA). Unpolymerized resin on the surface of newly fabricated bioreactors was removed with >99% isopropyl alcohol using the Form Wash station with a secondary washing step in a separate container containing fresh >99% isopropyl alcohol. This two-step washing process was necessary to ensure removal of all unpolymerized resin from the surface. Washed bioreactors were then UV-cured using 405nm light for 80 minutes at 60°C (Form Cure, Formlabs), soaked in 1X PBS (ThermoFisher Scientific, USA) for three days with daily PBS exchanges, and then sterilized using ethylene oxide gas (EOGas 4, Andersen Sterilizers, UK). The complete stirred bioreactor culture platform consisted of one or more 100 mm Petri dishes housing SBRs and 12×3 mm stir bars (CG-2003-17, Chemglass Life Sciences, USA). The dishes were placed on a 9-position magnetic stir plate (DuraMag, Chemglass Life Sciences) in either a single or stacked configuration (never exceeding three dishes) inside a standard cell culture incubator.

### Generation and maintenance of hPSCs

Five hPSC lines, summarized in Supplementary Table 1, were generated from skin biopsies via a Sendai virus-based reprogramming approach, as previously described.^40^ All hPSC lines were maintained on TC-plates (Corning, USA) coated with hESC-qualified Matrigel (Corning, USA) using mTeSR1 medium (StemCell Technologies, Canada), as previously described with minor modifications.^8^ hPSCs were passaged at 60% - 80% confluency, using an EDTA-based protocol, as previously described.^40^ Detached cell aggregates were replated at a ratio of 1/5 to 1/10 into growth factor reduced (GFR)-Matrigel coated 100 mm plates using mTeSR1 medium supplemented with 10 μM of the ROCK inhibitor Y-27632 (Tocris, USA). For RO differentiation, the hPSC clones were used between P14 -P22.

Two hPSC culture formats were used to generate OVs: adherent monolayer colonies and embryoid bodies (EB) aggregates. OV induction in adherent colonies was performed as previously described with minor modifications.^8, 41^ In brief, once hPSCs reached 20-30% confluency, defined as day 0 (D0), the medium was switched to the modified neural induction medium (NIM), consisting of DMEM/F12 (1:1; ThermoFisher Scientific, USA), 1% N2 supplement (ThermoFisher Scientific, USA), 1% MEM non-essential amino acids (MilliporeSigma, USA) and 5 mM Nicotinamide (Sigma-Aldrich, USA), with media changes every other day. After four days of differentiation in the modified NIM (D4), the medium was switched to NIM without nicotinamide. On D6, SBRs were placed in designated cultures (n=3 per cell line) with 30 mL of media and stirred at 300 RPM. On D10, when the colonies had formed neural tube-like structures, the medium was switched to retinal induction medium (RIM), consisting of DMEM/F12 (3:1), 2% B27 without vitamin A (ThermoFisher Scientific, USA), 1% MEM non-essential amino acids and 1% antibiotic-antimycotic solution (Gibco, USA). Cultures were maintained for an additional 8-10 days, with medium changes every other day. Between D18-20, OV-like structures were dislodged using a cell scraper, transferred to two ultra-low attachment 100 mm Petri dishes (SBIO, USA), and maintained as free-floating cultures in RIM medium supplemented with 20 ng/ml Insulin-like Growth Factor 1 (IGF1) (ThermoFisher Scientific, USA).

OV induction in EB cultures was carried out as previously described with minor modifications.^8^ In brief, hPSCs were cultured to 20-30% confluency to form OVs as above. After dissociation, hPSCs were resuspended in mTeSR1 with 10 μM Y-27632 and transferred into 100-mm polyHEMA (Millipore-Sigma, USA)-coated Petri dish for EB formation. NIM was supplied to the culture at a ratio of 3:1 and 1:1 at differentiation days D1 and D2, respectively. At D3, the media was switched to 100% NIM, with media changes every 2–3 days. EBs from 100-mm polyHEMA-coated dishes were plated onto 100-mm TC dishes coated with BD GFR Matrigel at D7 and cultured in NIM for 9 additional days with media changes every 2 days. On D15, SBRs were placed in designated culture dishes (n=3 per cell line) with 30 mL of media and stirred at 300 RPM. At D16, the media was switched to RIM. The media was changed every 2 days until OV-like structures appeared, typically emerging between D25 and D30. OV-like structures were dislodged using a cell scraper, transferred to two ultra-low attachment 100 mm Petri dishes, and maintained as free-floating cultures in the RIM medium supplemented with 20 ng/ml IGF1.

Quantification of OV yield and size was obtained 2-3 days after transferring to free-floating culture for both adherent and EB cultures.

### Immunofluorescence

Freshly dislodged OVs (100 μL samples) were collected, fixed in 4% paraformaldehyde for 1 hour at room temperature, and then washed 3 times in 1X PBS. OVs were then cryoprotected using a sucrose gradient (10–30%) overnight and embedded in M1 embedding matrix (ThermoScientific, USA) before sectioning. Ten-micron sections were obtained using an NX70 Cryostat (ThermoScientific, USA). Sections were dried on Superfrost Plus Micro-scope slides (Fisher Scientific, USA) and stored at −20 °C until use. For immunostaining, after 1-hour incubation in blocking solution (Tris-buffered saline with 0.1% Tween 20, 5% donkey serum, and 0.3% Triton, TBST) at room temperature, the sections were incubated overnight at 4°C with primary antibodies (Supplementary table 2) against CHX10 and ZO-1. After three 5-min washes in TBST, appropriate Alexa Fluor-conjugated secondary antibodies and DAPI were added for 1 hour at room temperature. Samples underwent a final wash (three 5-min washes in TBST) and excess TBST was removed before mounting. One drop of Fluoromount-G (Invitrogen, USA) was used to coat each sample before protecting with a coverslip and sealant.

The pimonidazole-based Hypoxyprobe-1 kit (Supplementary table 2) was used for immunostaining of hypoxic markers. Pimonidazole is a chemical compound reduced in hypoxic cells when the oxygen concentration is below 1.32% (10 mmHg pO_2_). Briefly, on D18 of adherent culture, 35 µL of pimonidazole HCL (200 µM) was added to 35 mL of culture media, and the samples were incubated for 16 hours in the incubator. Adherent OVs were then dislodged, fixed, cryoprotected, and sectioned as described above. After 1-hour incubation in blocking solution at room temperature, sections were incubated with primary antibody MAb1 (rat IgG1 monoclonal antibody) overnight, followed by Alexa Fluor-conjugated secondary antibody and DAPI. For apoptosis detection, the *in situ* Cell Death Detection Kit (TUNEL assay, Supplementary table 2) was used for immunostaining of apoptotic markers with samples processed exactly as described above. After fixation, sectioning, and blocking as described above, samples were incubated at 37°C for 1 hour with 50 µL of TUNEL reaction mixture per manufacturer’s instructions. Samples were then washed and stained with DAPI (as previously described).

### Image acquisition and Statistical Analysis

Bright-field images were acquired using an EVOS XL Core Cell Imaging System (ThermoFisher Scientific, USA). Fluorescence images were acquired using a confocal microscope (*Zeiss LSM 880)*. Images were processed and analyzed using ImageJ. TUNEL assay images were analyzed using Trainable Weka Segmentation (ImageJ plugin).^42^ Comparisons of static and SBR conditions were performed using an unpaired Student’s *t*-test with statistical significance defined as *p* < 0.05.

### Oxygen Analysis

In some experiments, before hPSC seeding, self-adhesive, sterilized oxygen sensor spots (PyroScience, Germany) were fixed to the bottom of 100mm TC Petri dishes. Measurements in control and SBR conditions were recorded as % air saturation via a fiber optic sensor cable attached to a FireSting-PRO meter (PyroScience, Germany). Data was logged using the PyroWorkbench software (PyroScience, Germany). Before data acquisition, a two-point calibration (0% and 100% air saturation) was performed according to the manufacturer’s instructions at 37 °C using a 50 mL solution of sodium sulfite (30 g/L) in distilled water to achieve 0% calibration and 50 mL of aerated distilled water (filtered air perfused for 1 hour) to achieve 100% calibration. Continuous dissolved oxygen measurements using the hPSC2 line were acquired at a 1 Hz sampling rate over 24 hours (n = 3, for control and SBR). Cyclical oxygenation measurements using the hPSC1 line were acquired at a 1 Hz sampling rate over 24 hours (n = 3, control and SBR). An intermittent agitation profile was created for cyclical oxygenation measurements using a two-outlet digital timer (Titan Controls, Canada), activating and deactivating the magnetic stir plate over three-hour intervals. Measurements were taken after each media change. Datasets were converted from % air saturation to % oxygen using PyroWorkbench software with media salinity set to 6.53 g/L.

### Numerical Modeling of SBR Fluid Dynamics & Validation

As cell culture media is considered an incompressible Newtonian fluid, the fluid flow within the system was modeled using COMSOL Multiphysics software (Stockholm, Sweden). Referencing the COMSOL application example model “Baffled Stirred Mixer” for component placement, the SBR and stir bar CAD designs were uploaded into the software. The simulation parameters were established by dividing the computational domain into a rotating inner region around the stir bar and a stationary outer region representing the rest of the Petri dish. A fine computational mesh (9.8 x 10^5^ tetrahedral volume elements with an average element quality of 0.67) was used to capture the details of the geometry, including additional refinement near the walls and the stir bar to ensure accurate modeling of the boundary layer effects. The rotation of the stir bar was set to 300 RPM. The simulation focused on laminar flow, excluding turbulence and gravity for simplicity. A time-dependent approach (t = 0 - 10 seconds) was used to simulate the motion, with time steps adjusted to accurately capture the rotation of the stir bar and its impact on the surrounding fluid with stir bar revolutions per minute equaling 300[1/min] * timestep(time[1/s]). The results of the simulation in terms of fluid velocity, vector positions, and shear stress were averaged to better approximate cumulative effects.

To validate the accuracy of the CFD simulation, particle image velocimetry (PIV) was used to visualize the fluid flow. Fluorescent polystyrene spheres (FluoSpheres (10 µm), Invitrogen, USA) were used as tracer particles to track particle paths and determine average velocities. Video recordings of sphere tracks were captured at 10 frames/sec using an Olympus BX50 Fluorescence Microscope (Evident, Japan) equipped with a high-speed CMOS camera (ORCA-Flash4.0 V3, Hamamatsu Photonics, Japan). Focusing on a plane of +2.5 mm from the bottom of the dish, an area of 3.34 mm^2^ was captured surrounding the exit aperture of one outlet port (Figure 6A). Focusing on this height allowed for a general characterization of fluid velocity flowing from the center reservoir into the outer domain of the SBR. Single frames of the video containing definitive sphere tracks were isolated, processed, and analyzed with ImageJ using the single-frame technique, wherein determining the displacement (ΔL) of the spheres between a known time interval (100 ms, Δt) allows for local fluid flow velocity (U) to be calculated (U= ΔL/Δt).

### RNA Extraction and RNA Sequencing Library Preparation

Two days after conversion from adherent to floating cultures, organoid samples were taken from the static and SBR cultured tissues. RNA sequencing data was generated from two hPSC lines (hPSC1 and hPSC2), following a protocol described previously.^8^ In brief, total RNA was isolated using the RNeasy Plus Mini Kit (Qiagen, USA), with each biological replicate comprising at least three retinal organoids from separate batches. The quality of the RNA was evaluated using the Bioanalyzer RNA 6000 Nano assay (Agilent, USA), and only samples with an RNA integrity number (RIN) greater than 8 were used for library construction. Then, 100 nanograms of total RNA were used to prepare strand-specific libraries with the TruSeq Stranded mRNA Library Prep Kit-v2 (Illumina, USA), incorporating slight modifications as described previously^43^. Samples were sequenced to an average depth of 14.5 million reads (range: 10.2 – 22.1 million reads) on a Illumina NextSeq 2000. Overall, 16 samples were analyzed, including two biological replicates from both static and SBR cultures for each hPSC line at day 20.

### Transcriptome Analysis

Reads were psuedoaligned to the human reference genome GRCH38.p7 with the Ensembl v106 annotation using kallisto v0.45.0. Transcript-level counts were summarized to the gene-level using tximport v1.30.0 then converted to counts per million and TMM normalized using edgeR v4.0.3. PCA and eigenvalue plots were generated using PCAtools v2.14.0. Upset plots were created with ComplexUpset v1.3.6. We performed GO term enrichment and created dotplots using clusterProfiler v4.10.0. Heatmaps were created with ComplexHeatmap v2.18.0 and single gene expression plots were generated with ggpubr v0.6.0.

---

**Supplementary Figure 1.**
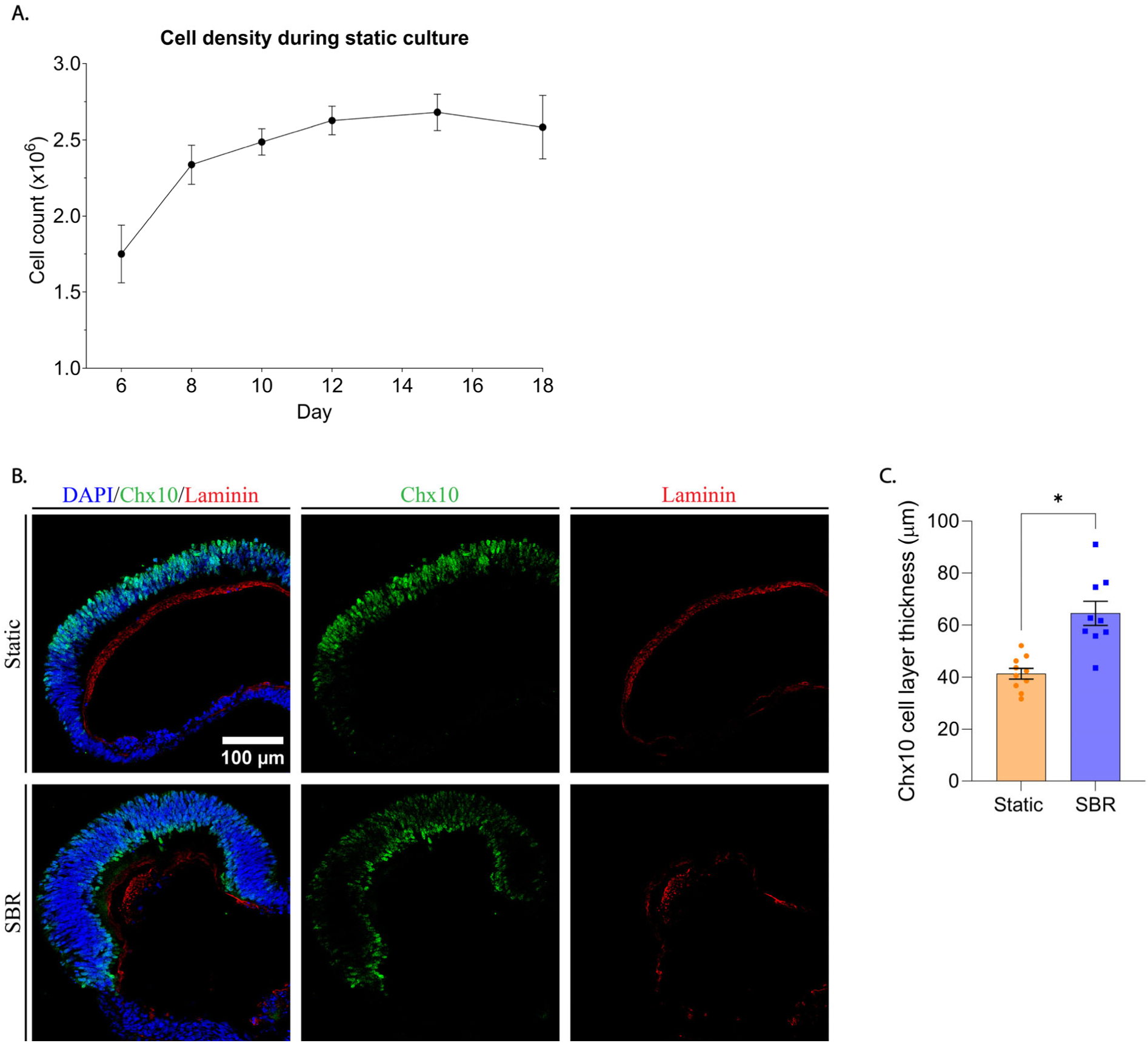
Cell Density During Static Culture and Chx10 Cell Layer Thickness. **<u>A.</u>** Analysis of cell density in static culture D6 to D18. **<u>B.</u>** Immunohistochemistry analysis of adherent OVs (hPSC1) using antibodies against markers for retinal progenitor cells or bipolar cells (CHX10, green), basement membrane (laminin, red). Nuclei were stained with 4′,6-diamidino-2-phenylindole (DAPI, blue). **<u>C.</u>** Quantification of the CHX10^+^ cell layer thickness of hPSC1 ROs produced by static and SBR conditions. The bar charts summarized data from ten measurements. Presented as mean ± standard deviation

**Supplementary Table 1.**
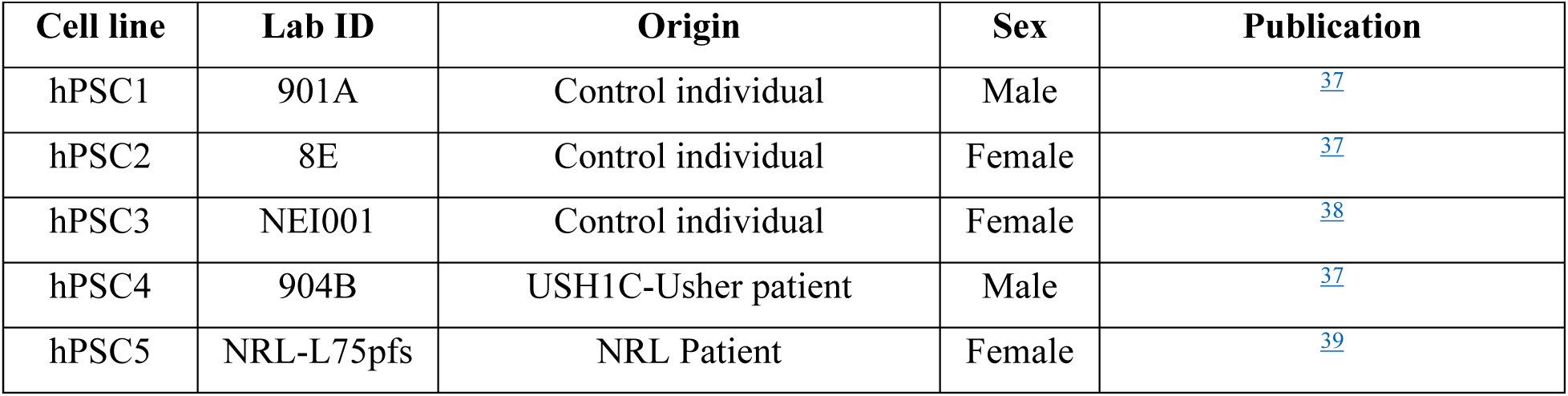
(hPSC Lines)

| Cell line | Lab ID | Origin | Sex | Publication |
| --- | --- | --- | --- | --- |
| hPSC1 | 901A | Control individual | Male | 37 |
| hPSC2 | 8E | Control individual | Female | 37 |
| hPSC3 | NEI001 | Control individual | Female | 38 |
| hPSC4 | 904B | USH1C-Usher patient | Male | 37 |
| hPSC5 | NRL-L75pfs | NRL Patient | Female | 39 |

**Supplementary Table 2.**
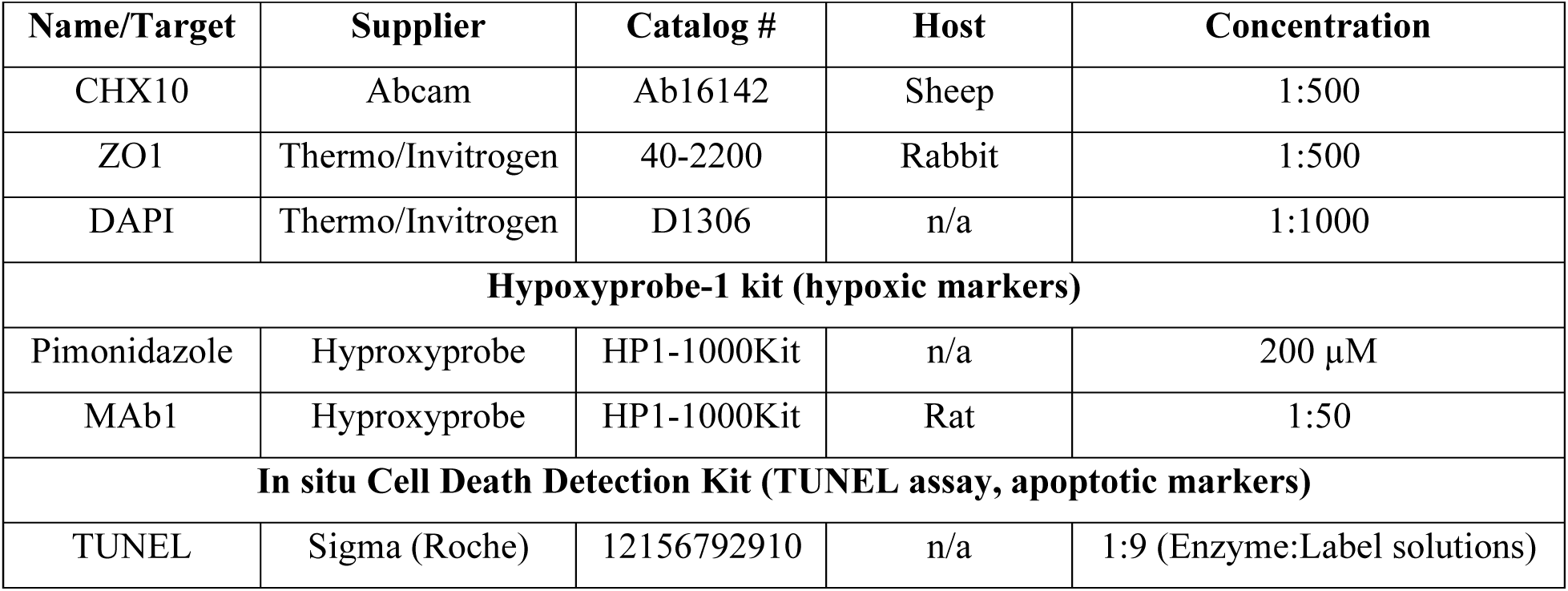
(Antibodies & Reagents)

**Supplementary Table 3.**
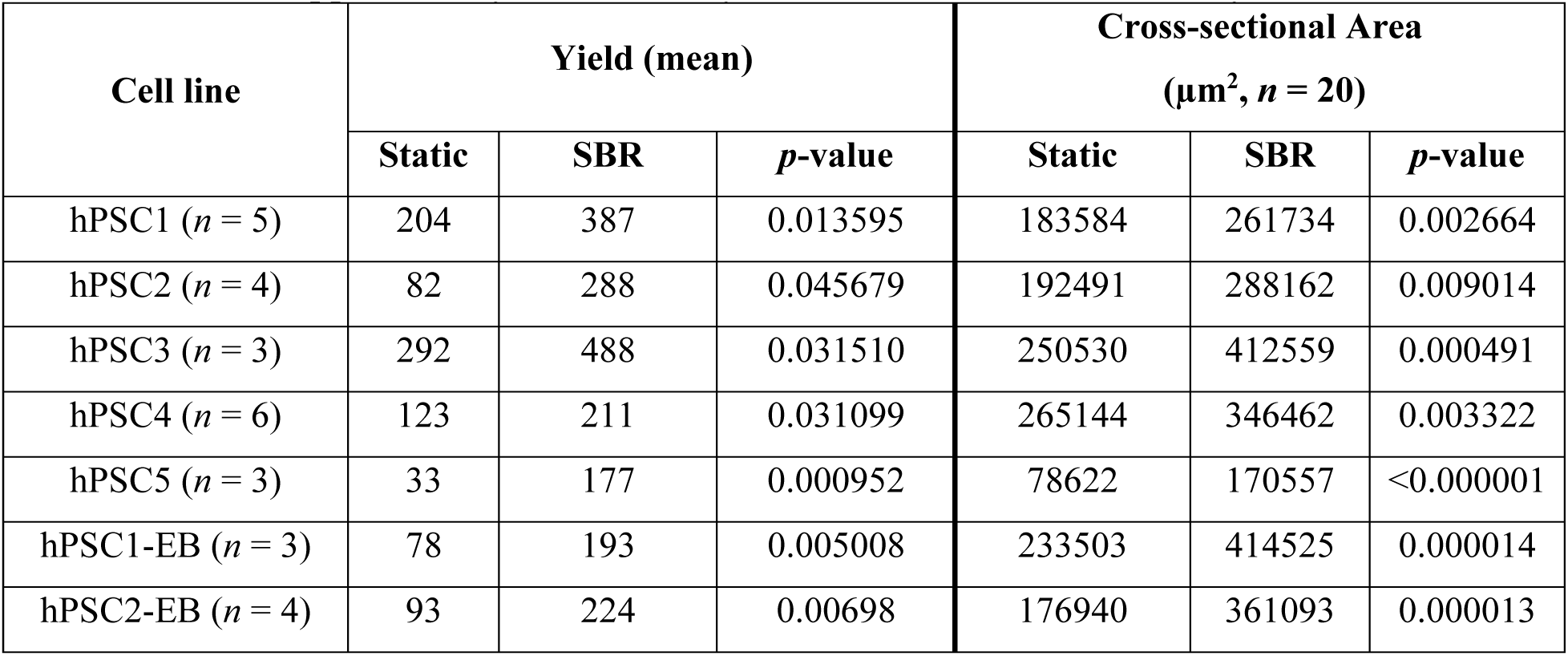
(OV yield & cross-sectional area analysis)

**Supplementary Table 4.**
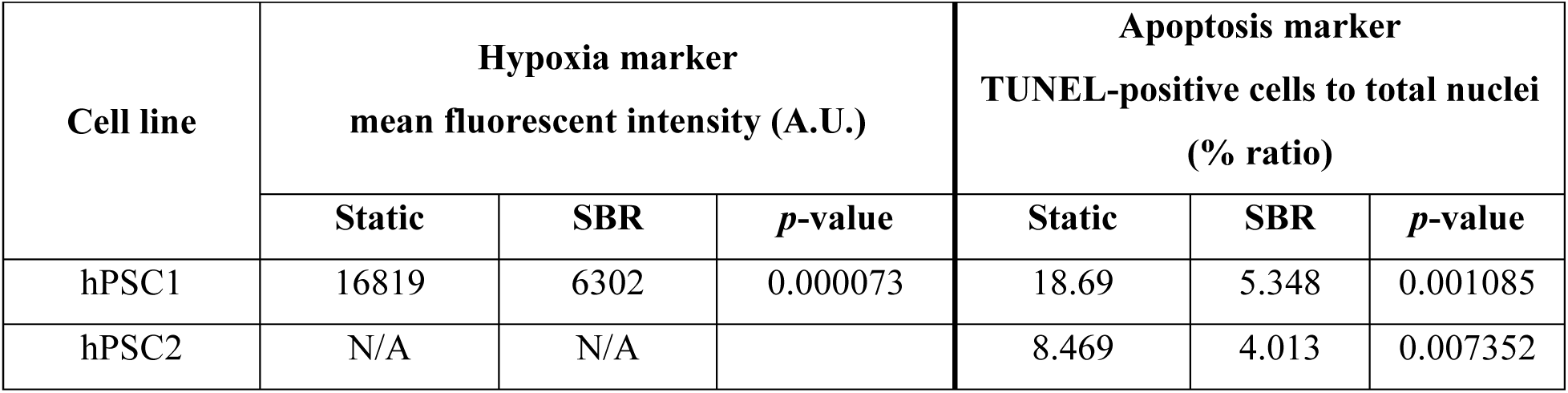
(Hypoxic and apoptotic IHC analysis)

**Supplementary Table 5.**
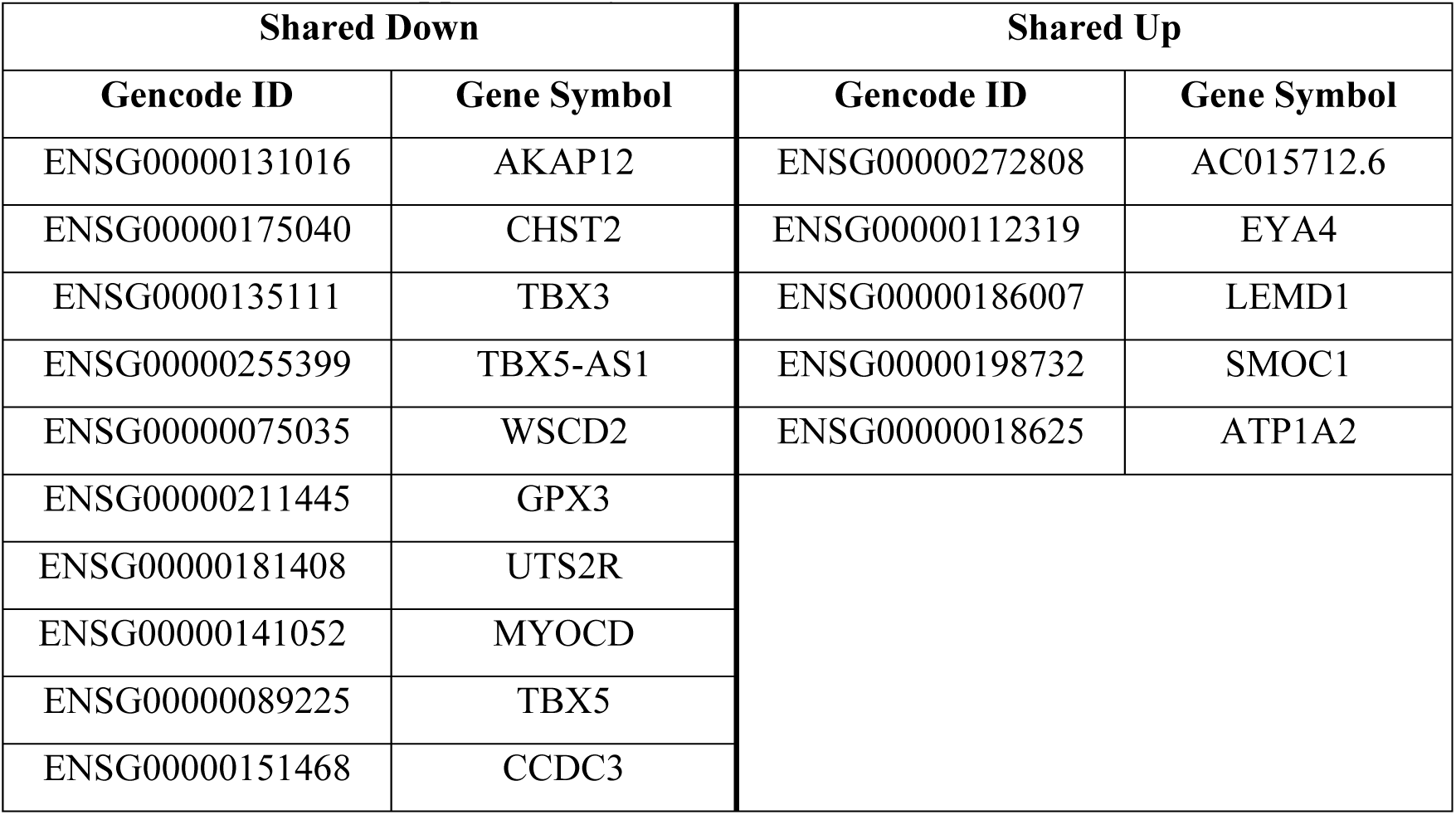
(Shared DE Genes)

