## Supplementary Tables for "Simple 3D-Printed Stirred Bioreactor Enhances Retinal Organoid Production Via Improved Oxygenation"

### Supplementary Table 1 (hPSC Lines)

| **Cell line** | **Lab ID** | **Origin** | **Sex** | **Publication** |
| --- | --- | --- | --- | --- |
| hPSC1 | 901A | Control individual | Male | ^41^ |
| hPSC2 | 8E | Control individual | Female | ^41^ |
| hPSC3 | NEI001 | Control individual | Female | ^42^ |
| hPSC4 | 904B | USH1C-Usher patient | Male | ^41^ |
| hPSC5 | NRL-L75pfs | NRL Patient | Female | ^43^ |

### Supplementary Table 2 (Antibodies & Reagents)

| **Name/Target** | **Supplier** | **Catalog #** | **Host** | **Concentration** |
| --- | --- | --- | --- | --- |
| CHX10 | Abcam | Ab16142 | Sheep | 1:500 |
| ZO1 | Thermo/Invitrogen | 40-2200 | Rabbit | 1:500 |
| DAPI | Thermo/Invitrogen | D1306 | n/a | 1:1000 |
| **Hypoxyprobe-1 kit (hypoxic markers)** | | | | |
| Pimonidazole | Hyproxyprobe | HP1-1000Kit | n/a | 200 µM |
| MAb1 | Hyproxyprobe | HP1-1000Kit | Rat | 1:50 |
| **In situ Cell Death Detection Kit (TUNEL assay, apoptotic markers)** | | | | |
| TUNEL | Sigma (Roche) | 12156792910 | n/a | 1:9 (Enzyme:Label solutions) |

### Supplementary Table 3 (OV yield & cross-sectional area analysis)

| **Cell line** | **Yield (mean)** | | | **Cross-sectional Area**  **(µm^2^, *n* = 20)** | | |
| --- | --- | --- | --- | --- | --- | --- |
|  | **Static** | **SBR** | ***p*-value** | **Static** | **SBR** | ***p*-value** |
| hPSC1 (*n* = 5) | 204 | 387 | 0.013595 | 183584 | 261734 | 0.002664 |
| hPSC2 (*n* = 4) | 82 | 288 | 0.045679 | 192491 | 288162 | 0.009014 |
| hPSC3 (*n* = 3) | 292 | 488 | 0.031510 | 250530 | 412559 | 0.000491 |
| hPSC4 (*n* = 6) | 123 | 211 | 0.031099 | 265144 | 346462 | 0.003322 |
| hPSC5 (*n* = 3) | 33 | 177 | 0.000952 | 78622 | 170557 | <0.000001 |
| hPSC1-EB (*n* = 3) | 78 | 193 | 0.005008 | 233503 | 414525 | 0.000014 |
| hPSC2-EB (*n* = 4) | 93 | 224 | 0.00698 | 176940 | 361093 | 0.000013 |

### Supplementary Table 4 (Hypoxic and apoptotic IHC analysis)

| **Cell line** | **Hypoxia marker**  **mean fluorescent intensity (A.U.)** | | | **Apoptosis marker**  **TUNEL-positive cells to total nuclei (% ratio)** | | |
| --- | --- | --- | --- | --- | --- | --- |
|  | **Static** | **SBR** | ***p*-value** | **Static** | **SBR** | ***p*-value** |
| hPSC1 | 16819 | 6302 | 0.000073 | 18.69 | 5.348 | 0.001085 |
| hPSC2 | N/A | N/A |  | 8.469 | 4.013 | 0.007352 |

### Supplementary Table 5 (Shared DE Genes)

| **Shared Down** | | **Shared Up** | |
| --- | --- | --- | --- |
| **Gencode ID** | **Gene Symbol** | **Gencode ID** | **Gene Symbol** |
| ENSG00000131016 | AKAP12 | ENSG00000272808 | AC015712.6 |
| ENSG00000175040 | CHST2 | ENSG00000112319 | EYA4 |
| ENSG0000135111 | TBX3 | ENSG00000186007 | LEMD1 |
| ENSG00000255399 | TBX5-AS1 | ENSG00000198732 | SMOC1 |
| ENSG00000075035 | WSCD2 | ENSG00000018625 | ATP1A2 |
| ENSG00000211445 | GPX3 |  | |
| ENSG00000181408 | UTS2R |  |  |
| ENSG00000141052 | MYOCD |  |  |
| ENSG00000089225 | TBX5 |  |  |
| ENSG00000151468 | CCDC3 |  |  |
